# Cancer/Testis Antigens Differentially Expressed in Prostate Cancer: Potential New Biomarkers and Targets for Immunotherapies

**DOI:** 10.1101/646869

**Authors:** Luciane T. Kagohara, Neil M. Carleton, Sayuri Takahashi, Takumi Shiraishi, Steven M. Mooney, Robert L. Vessella, Robert H. Getzenberg, Prakash Kulkarni, Robert W. Veltri

## Abstract

Current clinical tests for prostate cancer (PCa), such as the PSA test, are not fully capable of discerning patients that are highly likely to develop metastatic prostate cancer (MPCa). Hence, more accurate prediction tools are needed to provide treatment strategies that are focused on the different risk groups. Cancer/testis antigens (CTAs) are expressed during embryonic development and present aberrant expression in cancer making them ideal tumor specific biomarkers. Here, the potential use of a panel of CTAs as a biomarker for PCa detection as well as metastasis prediction is explored. We initially identified eight CTAs (*CEP55, NUF2, PAGE4, PBK, RQCD1, SPAG4, SSX2* and *TTK*) that are differentially expressed in MPCa when compared to local disease and used this panel to compare the gene and protein expression profiles in paired PCa and normal adjacent prostate tissue. We identified differential expression of all eight CTAs at the protein level when comparing 80 paired samples of PCa and the adjacent non-cancer tissue. Using multiple logistic regression we also show that a panel of these CTAs present high accuracy to discriminate normal from tumor samples. In summary, this study provides evidence that a panel of CTAs, differentially expressed in aggressive PCa, is a potential biomarker for diagnosis and prognosis to be used in combination with the current clinically available tools and is also a potential target for immunotherapy development.

## 1. Introduction

Prostate cancer (PCa) is the most prevalent cancer type among men and the second leading cause of male cancer-associated deaths in the United States accounting for an estimated 165,000 new cases and 30,000 deaths in 2018. While local tumors are successfully treated, metastatic PCa (MPCa) remains an incurable disease with a 30% 5-year survival rate [1]. The most common treatment for advanced PCa consists of androgen ablation to which most patients are responsive; however, a great proportion of men progress with metastatic castration-resistant PCa (mCRPC) and die from the disease. Although much progress in the treatment of mCRPC was made in the last decade, improvements regarding survival are still measured in months [2,3].

PCa screening and disease control is largely based on the prostate specific antigen (PSA) test that was initially introduced as a follow-up instrument for the detection of recurrence and progression to metastatic disease. Subsequently, its potential as an early diagnostic tool was explored [4,5] and PSA was accepted as a standard test to identify men at risk of PCa before any symptoms appeared. Thus, PSA was heralded as a promising early detection biomarker [5]. However, PSA screening has been considered a controversial assessment since many men are over-diagnosed and over-treated since PSA is not capable of differentiating more indolent from aggressive disease. It is estimated that 23% to 60% of men, with increased PSA levels present with prostate tumors that would remain clinically insignificant during their lifetime [6]. Unfortunately, these men who present with increased PSA may be submitted to unnecessary aggressive and invasive treatment and its consequent comorbidities [6–8] [6–8]. The use of active surveillance programs in men who are considered to have very low and low risk prostate cancer has had a major impact on over-treatment but one of the major dilemmas in PCa remains to identify patients with aggressive tumors at an early stage so that they can benefit from immediate definitive treatment. PSA based tests such as the Prostate Health Index (phi) and the 4K Score, are options to predict more accurately detect PCa [9]. The first test, that measures total, free and [-2]proPSA [10]; is FDA approved and have shown to be an important tool for risk stratification [11,12]. The 4K Score measures four kallikrein markers (total, free and intact PSA and hK2) and presents the same performance and is also associated with the risk of MPCa [13,14]. Still, additional molecular biomarkers for a combined test are crucial to categorize tumors according to their aggressive potential in a more accurate manner and to stratify men with PCa into more appropriate treatment strategies.

Cancer/testis antigens (CTAs) constitute an important class of cancer biomarkers that have not been fully explored, especially in PCa [15–18]. CTAs by definition are normally expressed in testis and other developmentally regulated tissues (e.g., placenta) but are aberrantly expressed in many types of cancers [19]. This unique pattern of expression makes these genes attractive candidates as biomarkers and, together with their immunogenic capacity, also good targets for the development of cancer immunotherapy [20–22]. The aberrant expression of CTAs in different cancer types is associated with phenotypic changes that confer cancer cells added advantages for proliferation and survival [23,24]. In a previous study, Takahashi et al. [25] evaluated the expression of 22 CTAs in localized (LPCa) and MPCa. Five of the CTAs (*CEP55, NUF2, PAGE4, PBK* and *SPAG4*) were differentially expressed between the two groups, suggesting that CTAs have the potential as biomarkers for differentiating aggressive PCa. However, since it was a retrospective study, the possibility of using these CTAs as predictors for MPCa could not be assessed.

In this study, we used the data generated by Takahashi et al. [25] to create a panel of CTA genes that are differentially expressed between LPCa and MPCa, and used this gene set to develop a panel of biomarkers for PCa screening. We hypothesize that using a panel of genes differentially expressed in advanced PCa early in the screening process would facilitate the early prediction of patients that will develop metastasis. In addition to Takahashi et al. analysis [25], we used a statistical multivariate logistic regression (MLR) model to identify with more stringency, a panel of potential CTA candidates as biomarkers for aggressive tumors. We found that, among the CTAs evaluated in the current study, *PAGE4* is down-regulated (undetectable) in 100% of MPCa cases. Thus, PAGE4 is a promising candidate to discriminate indolent from aggressive cases. Also, our results showed that the CTAs *CEP55, NUF2, PBK* and *TTK* were up-regulated in MPCa and their combined pattern of expression was capable of differentiating metastatic from non-metastatic tumors. Finally, we evaluated the expression of this CTA panel in normal and tumor paired tissues from PCa patients who were treated with radical prostatectomy to identify their potential as screening biomarkers. We observed significant variation in mRNA and protein expression levels of all these CTAs, suggesting that the changes in expression occur before metastasis development and could be used as early diagnostic and prognostic biomarkers.

## 2. Materials and Methods

### 2.1. Clinical Samples

Samples from clinically localized PCa (LPCa) (n=20) and soft tissue metastasis (MPCa) (n=20) were obtained at University of Washington from radical prostatectomies and autopsies, respectively. The age range of the patients with clinically LPCa was 48-75 years (median, 58 years) and a preoperative serum PSA median of 7.54 (ng/ml) (range, 2.4-64.0). The Gleason Score was: 6 (n=3), 7 (n=14), 8 (n=1) and 9 (n=2). Soft tissue metastasis were obtained from lymph node (n=8), liver (n=5), adrenal (n=1), bladder (n=1), kidney (n=1), lung (n=1) and pancreas (n=1). The specimens were used with the approval of the University of Washington Institutional Review Board. Complete demographic and clinical data are presented on **Supplementary Table 1**. Approximately 30 to 100mg of fresh tissue (with no dimension greater than 0.5cm) was collected and placed in RNAlater Solution (Ambion, Austin, TX). Samples were stored at 4°C for 1-7 days to allow solution to thoroughly penetrate the tissue and then maintained at −20°C until RNA extraction [25].

RNA samples from matched tumor and normal adjacent tissues were obtained from the Prostate Cancer Biorepository Network (PCBN). Using the standard operating procedure (SOP) protocols, as previously described in detail [26], RNA was isolated from 24 radical prostatectomy specimens. The grade and stage of each case are listed in **Supplementary Table 2**. Each case consisted of fresh-frozen tumor and benign tissues obtained at radical prostatectomy. Cancer samples were macro-dissected to ensure the presence of at 70% to 90% tumor cells.

The paired normal and PCa samples for immunohistochemistry assays were included in tissue microarrays (TMAs). The two TMAs included 80 unique prostate cancer patients representing different Gleason scores (3+3, 3+4, 4+3, and ≥8) with quadruplicates of cancer and cancer-adjacent normal areas. The detailed demographics of the total 80 cases stratified by Gleason scores are shown in **Supplementary Table 3**.

### 2.2. RNA isolation

RNA from 20 paired normal and PCa from PCBN were obtained using Trizol (Invitrogen). RNA quantification and integrity were assessed by Nanodrop and 2100 Bioanalyzer (Agilent Technologies). Additional information for PCBN SOPs can be found at http://www.prostatebiorepository.org/protocols.

### 2.3. Nanostring gene expression analysis

Nanostring nCounter Gene Expression Assay (NanoString Technologies, Seattle, WA) gene expression data were obtained previously for the LPCa and MPCa cohort [25]. The Nanostring approach was performed for 22 CTA genes (*CEP55, CSAG2, CTAG1B (NY-ESO-1), JARID1B, MAGEA1, MAGEA2, MAGEA6, MAGEA12, NOL4, NUF2, PAGE4, PBK, PLAC1, RQCD1, SEMG1, SPAG4, SSX2, SSX4, TMEFF2, TMEM108, TPTE* and *TTK*). The CTA genes were selected by mining publicly available microarray data from the Gene Expression Omnibus (http://www.ncbi.nlm.nih.gov/geo) in conjunction with our own data [27,28]. *ACTB* was used as the housekeeping gene for normalization.

### 2.4. qRT-PCR gene expression analysis

One microgram of total RNA was used for cDNA synthesis using the iScript cDNA Synthesis Kit (Bio-Rad Laboratories, Inc., Hercules, CA). The PCR reactions were performed with 0.2 µl of cDNA template in 25 µl of reaction mixture containing 12.5µl of iQ SYBR Green Supermix (Bio-Rad Laboratories, Inc.) and 0.25 µmol/L each primer. PCR reactions were subjected to hot start at 95°C for 3 minutes followed by 45 cycles of denaturation at 95°C for 10 seconds, annealing at 60°C for 30 seconds, and extension at 72°C for 30 seconds using the CFX96 Real-Time PCR Detection System (Bio-Rad Laboratories, Inc.). Analysis and fold differences were determined using the comparative threshold cycle method. *ACTB* was the housekeeping gene used for normalization. Primers’ sequences for the CTAs evaluated are shown in **Supplementary Table 4**.

### 2.5. Immunohistochemistry

The TMA slides were deparaffinized using xylene, and tissues were rehydrated in decreasing concentrations of ethanol (100%, 75%, 50%, and 25%; all vol/vol). Antigen retrieval was performed at controlled pH values under heat, followed by endogenous peroxidase inhibition using 0.3% hydrogen peroxidase. TMA slides were incubated for 1h at room temperature with a proprietary protein block, Protein Block Serum Free reagent (Dako). Primary antibody incubation was performed at 4°C overnight using the ideal dilution for each antibody (**Supplementary Table 5**). Primary antibody was washed with 1X PBS, and secondary antibody (1:200) was added to the slides and incubated for 1h at room temperature. Antigen localization was developed using 3,3′-diaminobenzidine chromogen. Tissue samples were counterstained in hematoxylin and dehydrated in ethanol and xylene.

For quantitative IHC (qIHC) analysis, slides were scanned using the Aperio Scanscope XT (Leica Biosystems) and the staining quantifications were performed using Aperio Imagescope v12.3 software (Leica Biosystems). Intensity and frequency of positive staining are determined by the pixel count of the delimited area selected for analysis. Intensity (different brown-staining shades) for a determined area is given as the total brown pixel count for that region. The frequency (area of positive staining) is given by the ratio of positive brown region and the total area selected for analysis (positive + negative area). Protein expression differences between the paired normal and tumor areas were compared using the Wilcoxon matched-pairs test. The average for all cores available from each patient for qIHC analysis was calculated, and the values were used to compare medians between the groups (tumor vs. benign). Protein expression (frequency or intensity) was considered significantly different for a *P* value ≤0.05.

### 2.6. Statistical analysis

Receiver Operator Characteristic (ROC) curves were used to identify CTAs with a high probability of accurately discriminating between localized and metastatic PCa or tumor and non-tumor cases. Gene expression changes were considered significant when AUC>0.7. Wilcoxon signed-rank or Mann-Whitney non-parametric test were used to compare CTA gene expression means between LPCa vs. MPCa and benign vs. tumor tissues, respectively. Gene expression differences were considered significant when *P* value ≤0.05. After the best individual genes were identified, the multivariate logistic regression (MLR) backward stepwise model was used to identify a CTA panel (with high specificity, sensitivity and significant AUC) capable of discriminating LPCa from MPCa or tumor from benign cases. All statistical analyses were performed using STATA version 13.

## 3. Results

### 3.1. Differential CTA gene expression in LPCa and MPCa

Nanostring is a digital multiplex approach in which multiple mRNAs can be absolutely quantified making the cDNA synthesis step unnecessary. Using this approach, Takahashi et al. [25] measured the expression of a panel of 22 CTA genes. All analyses were normalized using *ACTB* as a house-keeping gene. Here, we used the previously published dataset to perform a more stringent statistical analysis to identify CTAs that can accurately discriminate LPCa from MPCa.

We performed ROC analyses to verify the accuracy of each biomarker expression profile in discriminating LPCa from and MPCa samples. To classify the 22 CTA genes (*CEP55, CSAG2, CTAG1B (NY-ESO-1), JARID1B, MAGEA1, MAGEA2, MAGEA6, MAGEA12, NOL4, NUF2, PAGE4, PBK, PLAC1, RQCD1, SEMG1, SPAG4, SSX2, SSX4, TMEFF2, TMEM108, TPTE* and *TTK*) as good markers to discriminate indolent and aggressive cases, we used a cutoff AUC≥0.7. ROC curve analysis was also used to determine the highest specificity, sensitivity, positive (PPV) and negative prediction (NPV) values that maximize the cases correctly classified. Expression level means were compared to assure that the differences found were significant. Nanostring multiplex gene expression analysis of the CTA genes showed down-regulation of *PAGE4* and up-regulation of *CEP55, MAGEA2, NUF2, PBK, RQCD1, SPAG4, SSX2*, and *TTK* in MPCa (compared with LPCa) (Figure 1A, Table 1 and **Supplementary Figure 1**) with AUC above the cutoff established, suggesting that each of the CTAs was capable of discriminating the two groups. *PAGE4* was at undetectable levels in all MPCa cases.

**Table 1.** Localized and metastatic prostate cancer gene expression ROC analysis for 22 CTAs.

| CTA | NANOSTRING |  |  |  |  | qRT-PCR |  |  |  |  |
| --- | --- | --- | --- | --- | --- | --- | --- | --- | --- | --- |
|  | AUC | SENSITIVITY | SPECIFICITY | PPV | NPV | AUC | SENSITIVITY | SPECIFICITY | PPV | NPV |
| *PAGE4* | 0.99 | 95.00 | 95.00 | 95.00 | 95.00 | 0.99 | 95.00 | 90.00 | 90.48 | 94.74 |
| *CEP55* | 0.97 | 90.00 | 90.00 | 90.00 | 90.00 | 0.91 | 80.00 | 90.00 | 88.89 | 81.82 |
| *NUF2* | 0.92 | 80.00 | 95.00 | 94.12 | 82.61 | 0.89 | 70.00 | 90.00 | 87.50 | 75.00 |
| *PBK* | 0.86 | 65.00 | 90.00 | 86.67 | 72.00 | 0.81 | 55.00 | 85.00 | 78.57 | 65.38 |
| *SPAG4* | 0.85 | 70.00 | 80.00 | 77.78 | 72.73 | 0.80 | 65.00 | 95.00 | 92.86 | 73.08 |
| *TTK* | 0.81 | 65.00 | 85.00 | 81.25 | 70.83 | 0.76 | 50.00 | 90.00 | 83.33 | 64.29 |
| *RQCD1* | 0.79 | 65.00 | 85.00 | 81.25 | 70.83 | 0.79 | 60.00 | 85.00 | 80.00 | 68.00 |
| *SSX2* | 0.75 | 65.00 | 65.00 | 65.00 | 65.00 | 0.75 | 50.00 | 95.00 | 90.91 | 65.52 |
| *MAGEA2* | 0.71 | 45.00 | 80.00 | 69.23 | 59.26 | 0.63 | 40.00 | 65.00 | 53.33 | 52.00 |
| *SEMG1* | 0.70 | 80.00 | 45.00 | 59.26 | 69.23 | 0.52 | 35.00 | 70.00 | 53.85 | 51.85 |
| *TMEFF2* | 0.69 | 55.00 | 55.00 | 55.00 | 55.00 | 0.63 | 65.00 | 45.00 | 54.17 | 56.25 |
| *MAGEA6* | 0.69 | 50.00 | 85.00 | 76.92 | 62.96 | 0.68 | 40.00 | 85.00 | 72.73 | 58.62 |
| *MAGEA12* | 0.67 | 55.00 | 80.00 | 73.33 | 64.00 | 0.70 | 50.00 | 75.00 | 66.67 | 60.00 |
| *MAGEA1* | 0.67 | 50.00 | 85.00 | 76.92 | 62.96 | 0.75 | 45.00 | 80.00 | 69.23 | 59.26 |
| *CSAG2* | 0.63 | 40.00 | 80.00 | 66.67 | 57.14 | 0.72 | 50.00 | 80.00 | 71.43 | 61.54 |
| *PLAC1* | 0.57 | 45.00 | 75.00 | 64.29 | 57.69 | 0.59 | 45.00 | 75.00 | 64.29 | 57.69 |
| *CTAG1B* | 0.56 | 5.00 | 95.00 | 50.00 | 50.00 | 0.59 | 5.00 | 90.00 | 33.33 | 48.65 |
| *SSX4* | 0.51 | 40.00 | 65.00 | 53.33 | 52.00 | 0.80 | 55.00 | 85.00 | 78.57 | 65.38 |
| *JARID1B* | 0.51 | 45.00 | 45.00 | 45.00 | 45.00 | 0.67 | 50.00 | 85.00 | 76.92 | 62.96 |
| *TPTE* | 0.50 | 25.00 | 80.00 | 55.56 | 51.61 | 0.47 | 35.00 | 60.00 | 46.67 | 48.00 |
| *NOL4* | 0.46 | 25.00 | 70.00 | 45.45 | 48.28 | 0.55 | 35.00 | 75.00 | 58.33 | 53.57 |
| *TMEM108* | 0.42 | 40.00 | 60.00 | 50.00 | 50.00 | 0.68 | 40.00 | 75.00 | 61.54 | 55.56 |

CTAs: cancer/testis antigens; ROC: Receiver operating characteristic; AUC: area under curve; PPV: positive predictive value; NPV: negative predictive value.

**Figure 1.**
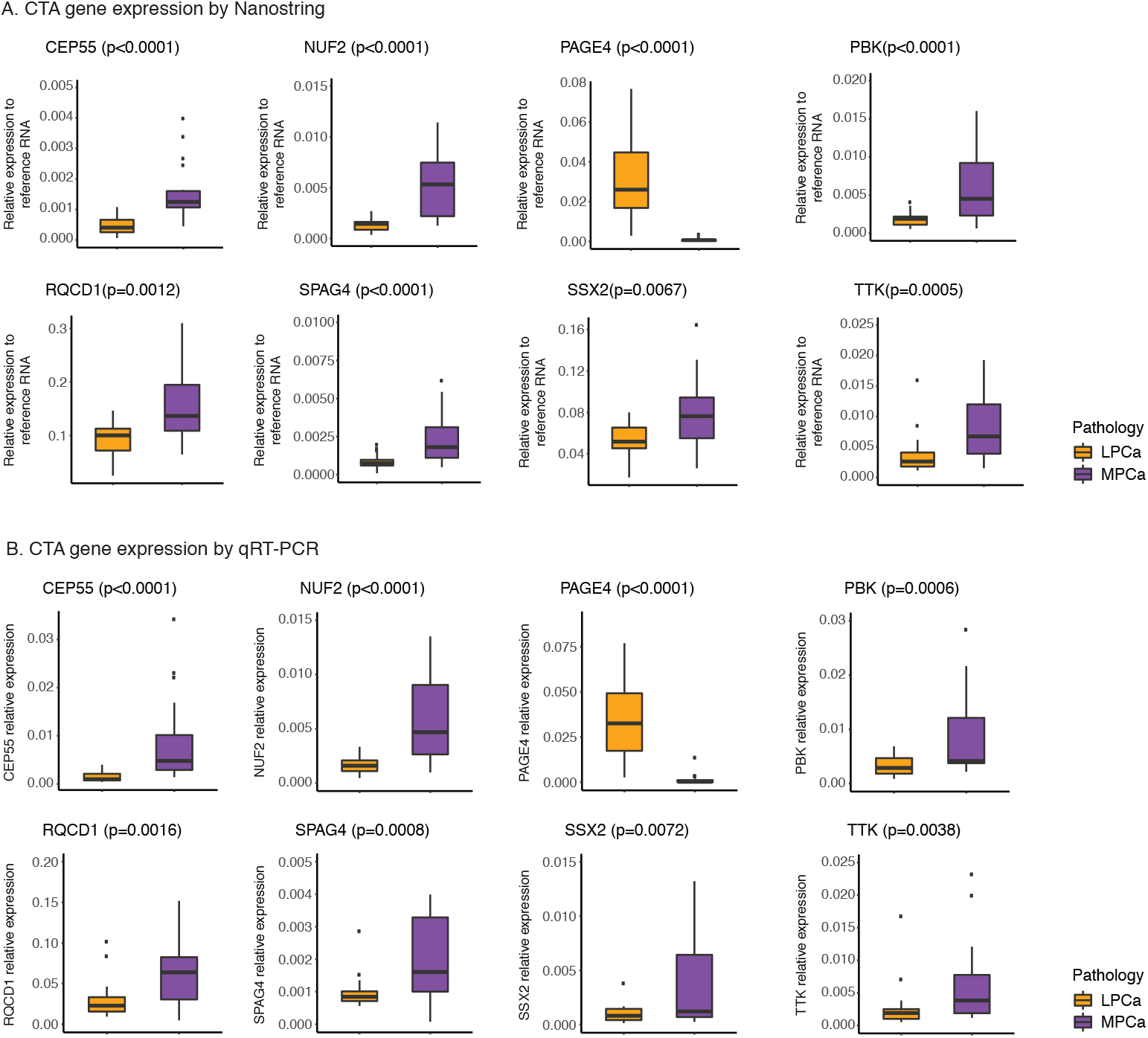
Cancer/testis antigens (CTA) gene expression analysis in localized (LPCa) and metastatic (MPCa) prostate cancer. Representation of gene expression measured by Nanostring (A) and by qRT-PCR (B). Nanostring relative gene expression is the ration between CTA and ActinB measured. For the qRT-PCR the relative gene expression calculation was performed using the 2^−ΔCt^ approach using ActinB as the housekeeping gene. Wilcoxon signed-rank test was used to compare means between LPCa and MPCa groups. Gene expression differences were considered significant when *P* value ≤0.05. *PAGE4* is down-regulated in MPCa while all other CTAs present increased expression. Nanostring results were confirmed by qRT-PCR in the same cohort (technical validation).

qRT-PCR was used to verify the results obtained using the Nanostring multiplex approach. Validation was performed for all 22 CTAs using the same sample sets that were examined by Takahashi et al [25]. Statistical analysis showed significant ROC curves (AUC>0.7) (Table 1) and confirmed overexpression of the CTA genes *CEP55, NUF2, PBK, RQCD1, SPAG4, SSX2* and *TTK* in MPCa, as well as the down-regulation of *PAGE4* (**Supplementary Figure 2** and Figure 1B). The other selected CTAs did not show significant expression changes between LPCa and MPCa (data not shown). Of note, in the study by Takahashi et al. [25], only *CEP55, NUF2, PBK, PAGE4* and *SPAG4* were found differentially expressed in LPCa vs. MPCa. However, in the present study, a more robust analysis increased the panel of potential aggressive PCa biomarkers. These data not only support the fact that CTA expression patterns can be used to discriminate MPCa and LPCa cases, but also corroborates the previous data using the same biomarkers.

**Figure 2.**
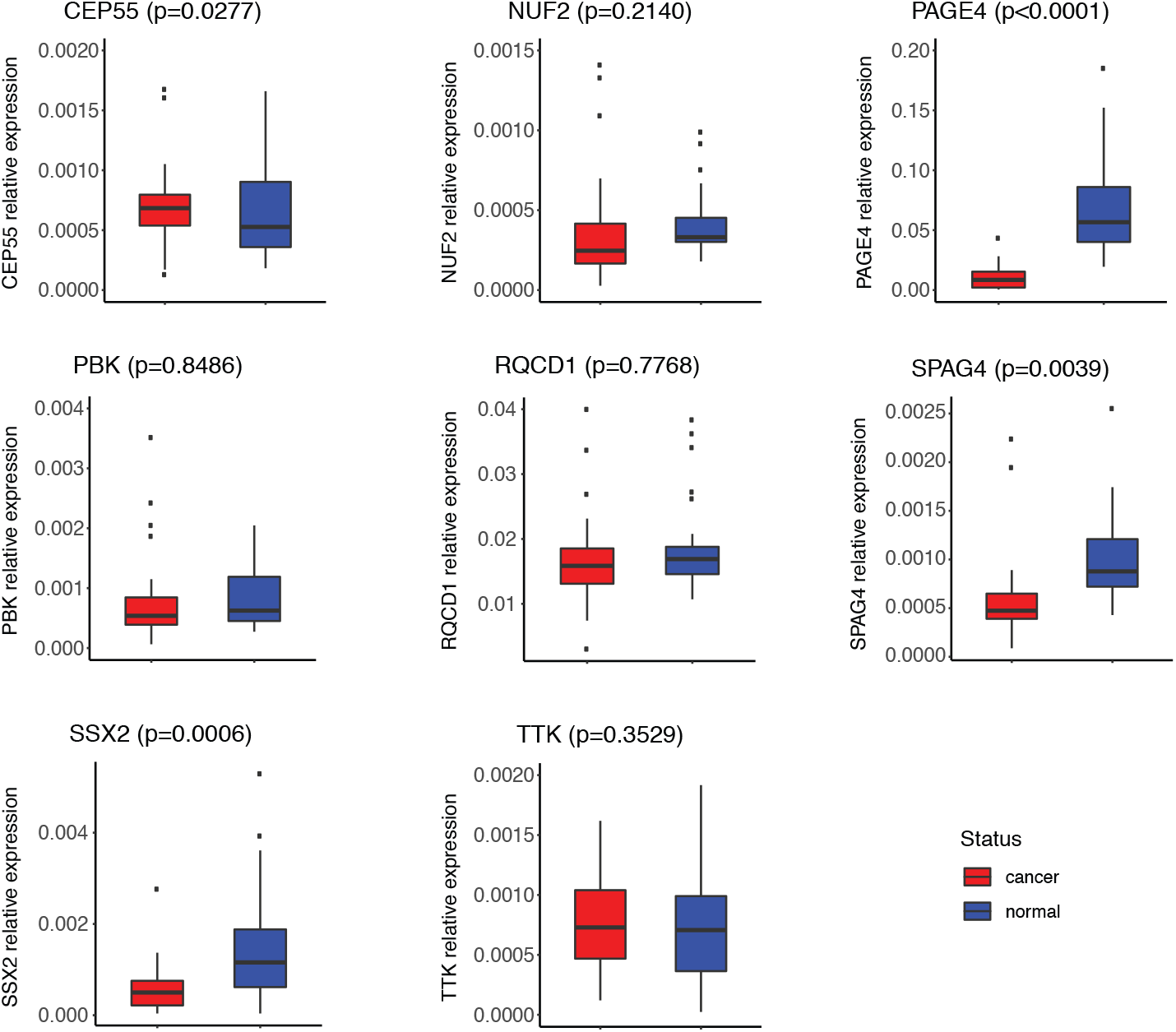
Cancer/testis antigens (CTA) gene expression analysis in paired tumor and normal adjacent samples from patients with prostate cancer. Gene expression was quantified by qRT-PCR. The relative gene expression calculation was performed using the 2^−ΔCt^ approach using ActinB as the housekeeping gene. Mann-Whitney non-parametric test was used to compare means between LPCa and MPCa groups. Gene expression differences were considered significant when *P* value ≤0.05. *CEP55* presents increased mRNA levels in PCa compared with normal samples. Up-regulation in normal versus tumor tissue was observed for *PAGE4, SPAG4* and *SSX2*. For the other CTAs no significant changes in expression was noted.

### 3.2. CTA expression in paired tumor and adjacent normal prostate tissue samples reveals differences at the mRNA and protein level

To determine if the CTAs differentially expressed in LPCa vs. MPCa also present different expression patterns in normal prostate tissue and PCa samples both at the mRNA and protein level, *CEP55, NUF2, PAGE4, PBK, RQCD1, SPAG4, SSX2* and *TTK* expression levels were evaluated in paired tumor samples and the adjacent normal tissues obtained from radical prostatectomies. Two distinct cohorts were used, one for gene expression analysis (22 paired samples) and another for protein expression (80 paired samples).

Gene expression analysis of the 22 paired tumor and normal samples did not show significant differences for *CEP55, NUF2, PBK, RQCD1* and *TTK* (Figure 2). *PAGE4, SPAG4* and *SSX2* are up-regulated in benign areas of the prostate when compared to tumor tissue. The expression profile of these genes can discriminate with good accuracy normal from PCa samples, as shown by ROC curve analysis (Table 2). These findings suggest that, for the CTAs selected in this study, changes in gene expression occur in advanced stages of PCa progression and are associated with a more aggressive phenotype.

**Table 2.**
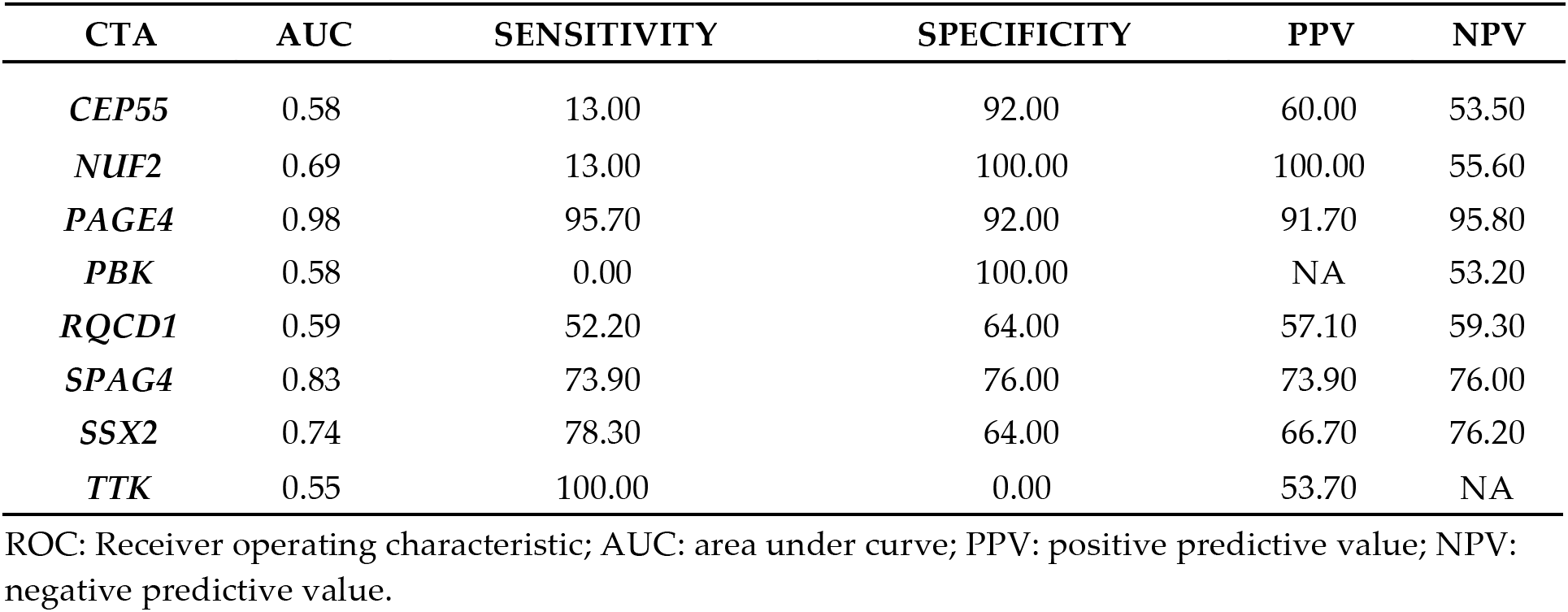
ROC analysis for gene expression profile of paired normal and tumor samples.

Although gene expression changes were not detected for some of the CTA genes selected, the protein expression analysis by IHC in 80 paired PCa and normal samples revealed significant differences between tumor and normal adjacent tissue from the same patients for all 8 genes (Figure 3A). Using quantitative image analysis, we measured the frequency and the intensity of the staining, independently. For both variables, we observed significant differences in the protein levels when comparing the tumor with its paired adjacent normal tissue (Mann-Whitney non-parametric test with *P* value ≤ 0.05) (Figure 3B and C). All CTAs show increased protein expression in PCa versus the normal tissue. In order to verify if the increased protein levels were useful to accurately discriminate tumor from adjacent normal samples we performed ROC analysis. The intensity of staining for all CTAS, but NUF2, is an accurate variable (AUC > 0.70) to discriminate cancer from normal tissue. (Table 3). The frequency of positively stained cells was significantly higher among tumor samples when compared to the normal adjacent paired tissue for all CTAs. Although, almost all AUCs were below the cutoff value (Table 3), when we compared the means of positive cells between tumor and normal samples the differences are significant (Figure 3B). The progressive down-regulation of *PAGE4* in PCa is a distinctive marker of metastasis development and therefore, tracing its loss of expression from the time of PCa diagnosis can provide valuable prognostic information. The observation that gene expression is not different between normal versus PCa and the fact that *SPAG4* and *SSX2* present with higher levels in normal samples suggest that slight changes at the transcriptional level may lead to significant changes in protein expression in cancer cells.

**Table 3.** ROC analysis summary for the immunohistochemistry intensity and frequency protein expression analysis.

| CTA | INTENSITY |  |  |  |  | FREQUENCY |  |  |  |  |
| --- | --- | --- | --- | --- | --- | --- | --- | --- | --- | --- |
|  | AUC | SENSITIVITY | SPECIFICITY | PPV | NPV | AUC | SENSITIVITY | SPECIFICITY | PPV | NPV |
| *CEP55* | 0.79 | 0.64 | 0.75 | 0.71 | 0.68 | 0.71 | 0.70 | 0.62 | 0.64 | 0.68 |
| *NUF2* | 0.67 | 0.54 | 0.73 | 0.66 | 0.63 | 0.69 | 0.75 | 0.53 | 0.61 | 0.69 |
| *PAGE4* | 0.72 | 0.56 | 0.78 | 0.71 | 0.65 | 0.61 | 0.64 | 0.54 | 0.57 | 0.62 |
| *PBK* | 0.78 | 0.65 | 0.82 | 0.77 | 0.72 | 0.64 | 0.71 | 0.47 | 0.56 | 0.64 |
| *RQCD1* | 0.71 | 0.63 | 0.73 | 0.69 | 0.66 | 0.64 | 0.71 | 0.52 | 0.59 | 0.64 |
| *SPAG4* | 0.72 | 0.61 | 0.77 | 0.72 | 0.67 | 0.62 | 0.69 | 0.49 | 0.57 | 0.62 |
| *SSX2* | 0.73 | 0.56 | 0.80 | 0.72 | 0.66 | 0.67 | 0.67 | 0.56 | 0.58 | 0.64 |
| *TTK* | 0.73 | 0.58 | 0.76 | 0.70 | 0.65 | 0.67 | 0.67 | 0.58 | 0.61 | 0.64 |

ROC: Receiver operating characteristic; AUC: area under curve; PPV: positive predictive value; NPV: negative predictive value.

**Figure 3.**
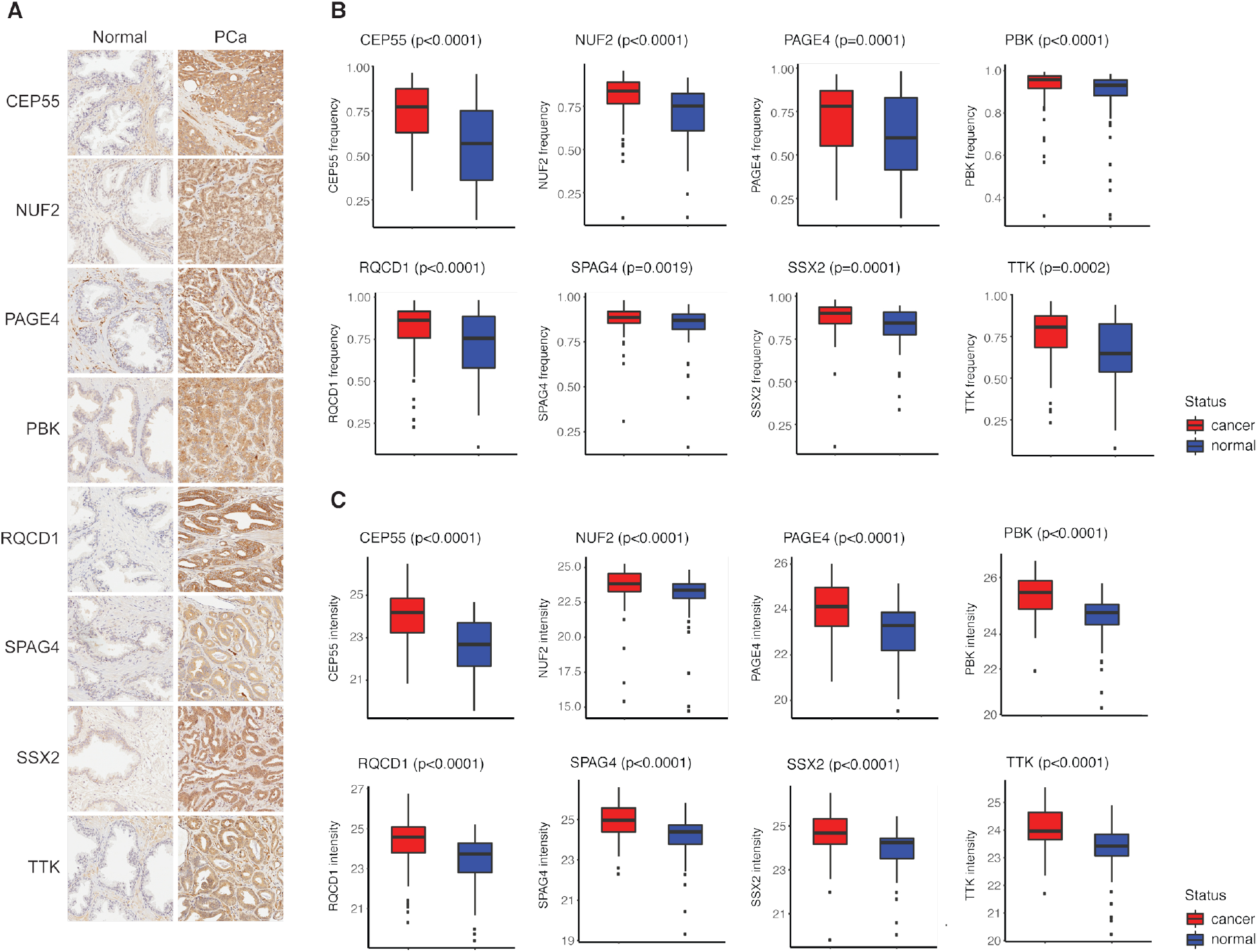
Cancer/testis antigens (CTA) protein expression analysis by immunohistochemistry (IHC) in paired tumor and normal adjacent tissues. IHC using antibodies against eight CTAs were performed to identify significant differences between normal and tumor areas from the same prostate. Panel A represents the immuno-staining for CTAs in normal and PCa paired samples. Using a computational quantitative approach, it was possible to measure the frequency (B) and intensity (C) of the staining. Mann-Whitney non-parametric test was used to compare means (*P* value ≤0.05). All CTA proteins present increased expression in PCa when compared to the normal paired sample.

### 3.3. Identification of a CTA panel as a potential biomarker for aggressive disease

In an attempt to identify a panel of CTAs that would be more sensitive than a single CTA to discriminate indolent from aggressive disease, we performed multiple logistic regression (MLR). We identified a panel of CTAs whose combined expression pattern could represent a potential tool for the discrimination of MPCa cases from LPCa. Using the expression profiles determined by qRT-PCR, MLR led us to a panel that included the CTAs *CEP55* and *RQCD1* and that correctly classify MPCa or LPCa in 87.5% of the cases evaluated in the present study (AUC=0.95, sensitivity=85.0%; specificity=90.0%; positive predictive value=89.5%) (Figure 4A).

**Figure 4.**
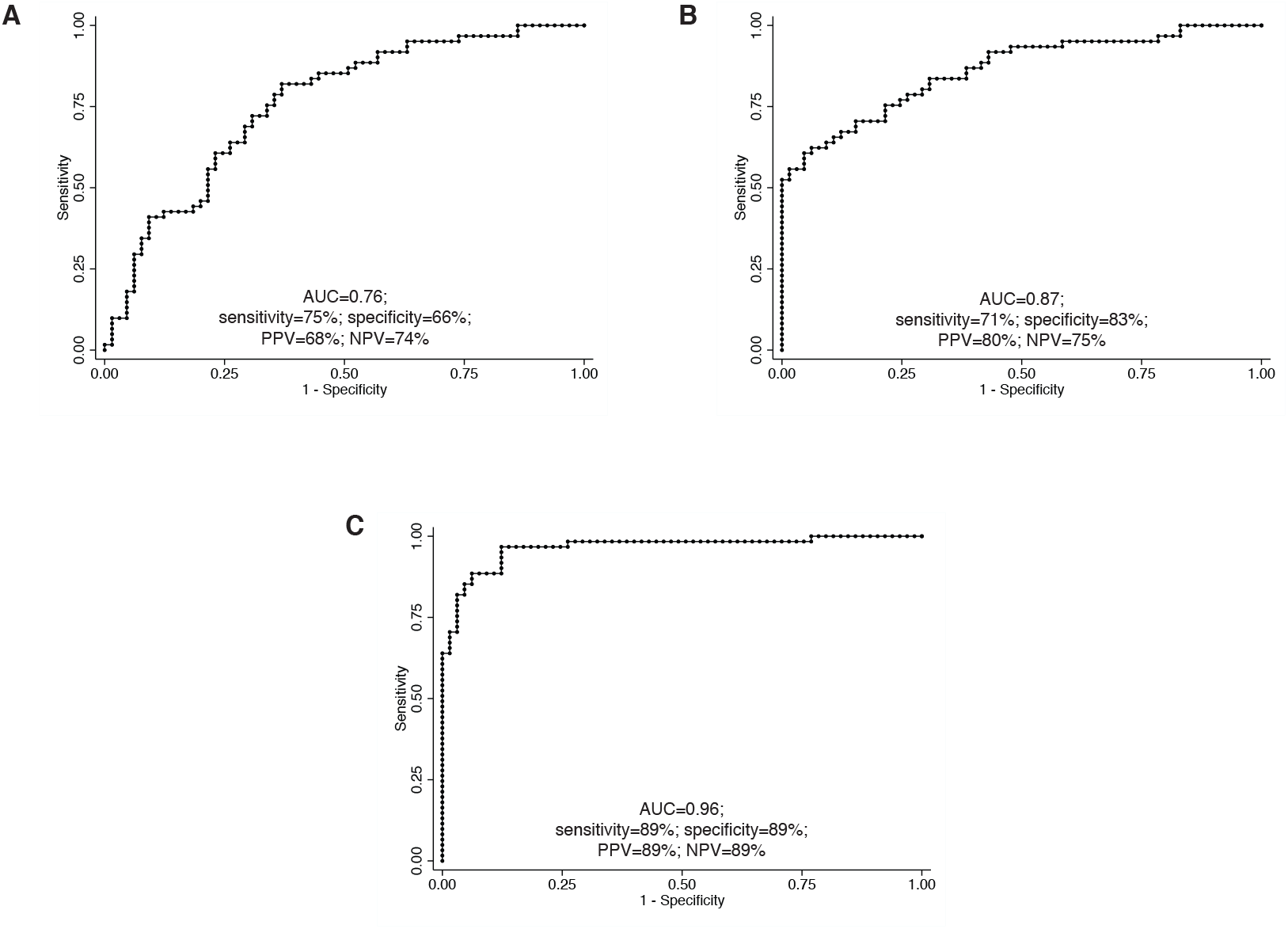
Receiver operating characteristic (ROC) curve analysis of the multivariate logistic regression (MLR) performed to identify panels of biomarkers accurate to discriminate localized (LPCa) from metastatic (MPCa) prostate cancer and normal from tumor samples. A. ROC curve analysis for the gene expression levels of *CEP55* and *RQCD1* to discriminate LPCa from MPCa. B. ROC curve analysis for the protein expression analysis. Here, MLR identified a panel including all CTAs staining intensity and 3 CTAs staining frequency as a good panel to discriminate normal from tumor samples. C. ROC curve analysis for *PAGE4* gene expression that alone is capable of discriminating virtually all MPCa cases from LPCa. AUC – area under curve; PPV – positive predictive value; NPV – negative predictive value.

For the paired PCa and normal adjacent tissue cohort, the MLR analysis resulted in a panel in which intensity and frequency of the CTA proteins accurately discriminate normal from tumor samples. The panel including all CTA protein expression intensity and NUF2, PBK, SSX2 and TTK protein expression frequency correctly classified ~89% of the samples (AUC=0.96, sensitivity=88.5%; specificity=89.2%; positive predictive value=89.2%) (Figure 4B).

Although the combined expression of *CEP55* and *RQCD1* expression presented high accuracy, this panel is not any more accurate than the *PAGE4* pattern of expression alone. *PAGE4* by itself is capable of separating almost all cases (AUC~1) (Figure 4C). No other CTA selected from the current study demonstrates the same degree of accuracy to differentiate MPCa and tumor cases from LPCa like PAGE4. Therefore, a decrease in PAGE4 level is an important, and to the best of our knowledge, a unique feature among CTAs known to be expressed in PCa, mainly in MPCa. Therefore, PAGE4 is a strong candidate as a biomarker for aggressive prostate tumors.

### 3.4. CTA expression and association with Gleason score

The Gleason score is an important feature considered to determine therapy and prognosis of PCa patients. Due to its relevance, we investigated the CTAs protein expression in 3+3/3+4 and 4+3/≥8 Gleason score groups. Although, 3+4 and 4+3 are score 7, it is widely known that the prognosis for these groups of men are significantly different and so we decided to group the first with more indolent tumors (3+3/3+4) and the later with the more aggressive (4+3/higher). Most of the CTAs analyzed by IHC in this study present increased protein levels in patients with higher Gleason scores when compared to lower scores (Figure 5A and B). Frequency of PAGE4, PBK and RQCD1 positive cells are significantly increased in PCa with Gleason score 4+3/higher (Figure 5A). When considering the intensity of the IHC staining, PAGE4, PBK, SPAG4 and SSX2 presented significant stronger staining associated with more aggressive histopathology (Figure 5B). These findings suggest that PAGE4 and PBK could be used to determine PCa prognosis since the frequency and protein levels together in the tumor cells present positive association with the Gleason score.

**Figure 5.**
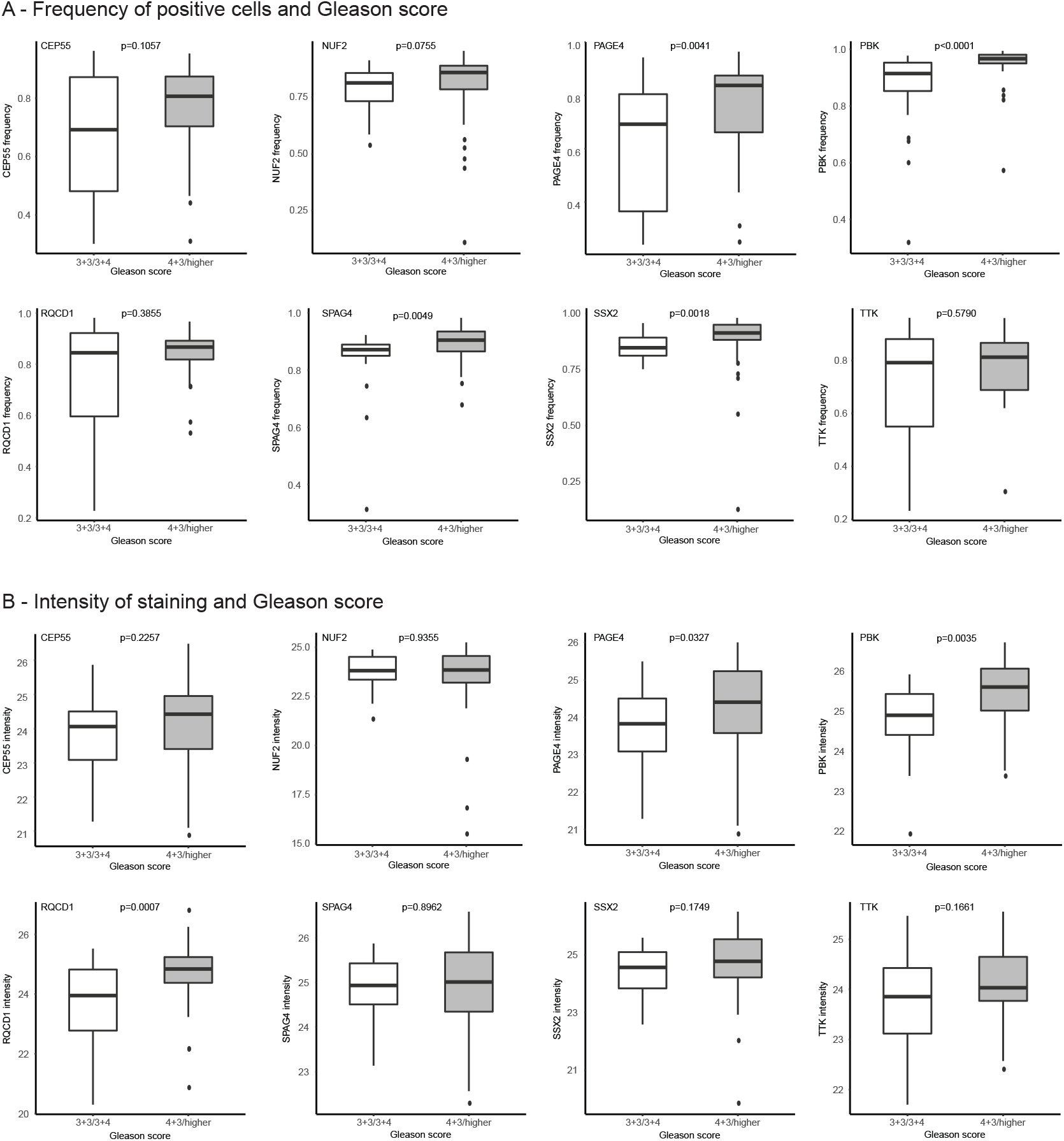
Cancer/testis antigens (CTA) protein expression by immunohistochemistry according to the Gleason score. The PCa samples from the TMAs were grouped in Gleason score 3+3/3+4 and 4+3/higher. The frequency of positive tumor cells (A) and the intensity of the staining (B) were measured in separate and statistical analysis (Mann-Whitnney non-parametric test) was performed for each of the staining measurements.

## 4. Discussion

In this study we used gene and protein expression analysis to identify a panel of CTA biomarkers that are differentially expressed in MPCa but could potentially be used for tumor screening as well. We used the gene expression profiles previously published by Takahashi et al. [25] and performed a new statistical exploration, with ROC curve and MLR analysis, to identify CTA genes that by themselves were capable of discriminating MPCa from LPCa cases and also to create a panel with even better accuracy. The expression of the CTAs *CEP55, NUF2, PAGE4, PBK, RQCD1, SPAG4, SSX2* and *TTK* are capable of discriminating aggressive from indolent PCa. Loss of *PAGE4* expression was detected in all MPCa cases and this feature alone is enough to distinguish all MPCa from LPCa cases. The up-regulation of *CEP55* and *RQCD1* together is the second most accurate biomarker of MPCa cases. These findings represent good evidence that the changes in CTA gene expression with the progression of PCa can identify men with more aggressive tumors. Unfortunately, since the LPCa and MPCa samples were not paired, it was not possible to determine the evolution of the CTA expression profiles in the same patient. Prospective studies with patient follow-up, from disease diagnosis until metastasis development, would allow a better understanding about the time point where the changes in CTA expression begin during the course of PCa development.

One of the main causes of death among men with PCa is metastatic disease [1]. Although PSA is the gold standard for screening, it lacks the ability to predict the development of metastasis [29,30]. Since the CTAs we selected from the Takahashi et al. [25] study are differentially expressed between LPCa and MPCa, we analyzed their expression in a cohort of paired tumor and adjacent normal tissue obtained from patients with PCa that underwent radical prostatectomy. Using quantitative IHC, we found that all the selected CTA proteins are up-regulated (intensity and frequency) in the tumor samples when compared to the normal adjacent tissue. As with the above mentioned cohort, we found a panel of CTAs whose intensity and frequency at the protein level are capable of discriminating normal from cancer samples with great accuracy. This observation further highlights the usefulness of CTAs as biomarkers for PCa that could be used during screening together with the PSA test, though further studies with larger cohorts across institutions and demographics are needed to determine the real prediction ability of these biomarkers for the development of MPCa. Studies in the future with prospective cohorts from screening to diagnosis, and the development of metastatic disease, can shed new light on how the expression of these CTAs progress during the course of the cancer.

An important contribution of this study is that we show the importance of using a panel of biomarkers for detection or prognosis. In both scenarios, normal vs. cancer, and LPCa vs. MPCa, the strongest predictors were those including more than one CTA. It is widely known that PCa and many other tumors are heterogeneous and composed of different cell populations with unique molecular profiles [31–34]. The use of single biomarkers may not cover the wide range of cell subclones present in the tumor and only capture the most abundant population. On the other hand, a panel of biomarkers is more likely to cover more broadly the different molecular profiles and allow the development of more accurate tests for screening and follow-up [35–37]. In the current study, there is one exception to our biomarker panel hypothesis: namely *PAGE4*. *PAGE4* gene expression was capable of discriminating MPCa from LPCa with 100% accuracy. Metastatic samples from men previously treated for PCa showed loss of this CTA expression compared to patients with local disease at the moment of diagnosis, suggesting that this gene is critical for tumor development but not for the metastasis establishment in a distant sites. This assumption is corroborated by the fact that PAGE4 protein expression is downregulated in metastatic PCa suggesting that PAGE4 may actually be a metastasis suppressor [38].

Recent studies suggest that *PAGE4* is developmentally regulated with dynamic expression patterns in the fetal prostate and that it is also a stress-response protein that is up-regulated in response to cellular stress [38]. In the present study, we observed loss of expression of *PAGE4* in in MPCa cases. Sampson et al. [39] also observed reduced levels of PAGE4 in MPCa when compared to indolent cases and found that in PAGE4 positive cells wild-type AR activity is reduced. This suggests that *PAGE4* plays an important role in MPCa, since aberrant activation of the AR pathway is a critical step in the progression to mCRPC after androgen ablation therapy. Loss of *PAGE4* in MPCa might result in activation of the AR signaling pathway, resulting in resistance to androgen-derivation therapy in men with MPCa [40,41]. However, as we recently demonstrated the role of PAGE4 in mCRPC is dependent on intratumor heterogeneity and downregulating the activity of the AR pathway depends on the co-expression and PAGE4 phosphorylation by HIPK1 and CLK2 that either potentiate or attenuate the effects on the AR pathway, respectively [42].

CTA expression in PCa and the association with disease aggressiveness was assessed previously in the same cohort of LPCa and MPCa by Takahashi et al. [25]. The authors found that *CEP55, NUF2, PAGE4, PBK* and *SPAG4* are differentially expressed in LPCa vs. MPCa. Using the same cohort, we repeated the technical validation and performed new statistical analysis using different tests. Besides the five CTAs previously shown to be differentially expressed between the two groups, we also found that *RQCD1, SSX2* and *TTK* are up-regulated in the MPCa samples. In addition, one of these CTAs, not previously predicted to be a biomarker candidate, was relevant when combined with another gene. The combined expression pattern of *CEP55* and *RQCD1* could be a marker for aggressive tumors. Although this panel is not any more accurate than *PAGE4* expression profile alone in local and metastatic tumors, it provides good evidence that a combination panel could be more relevant for prognostication than a single marker, resulting in higher specificity. Besides *PAGE4*, other CTAs such as *PBK* and *SSX2*, were previously detected in PCa. PBK expression is absent in vitro and in normal prostate tissue. A gradual increase in PBK expression is concurrent with increased disease aggressiveness [43], in accordance with our findings that this CTA is a marker of MPCa. *SSX2* expression was also detected in PCa samples in a few studies [44–46]. Smith et al. observed that SSX2 higher levels were present in advanced cases, however they also noticed that the pattern of expression across different tumor stages (including benign prostate) was heterogeneous [44]. The immunohistochemistry data and gene expression findings by Bloom & McNeel [47] corroborate our observations that SSX2 protein is increased in MPCa. The authors also showed that circulating tumor cells expressing the correspondent gene could only be detected in peripheral blood of PCa patients, while undetectable in healthy men [47].

One intriguing observation in our study is that the gene expression levels of *PAGE4, NUF2* and *SPAG4* are increased in normal prostate tissue relative to the tumor samples. Since the normal tissues were collected adjacent to the prostate tumors it is probable that the up-regulation of CTAs in non-cancer areas is a field effect as previously described by Zeng et al. [48]. They describe the same trend for *PAGE4* when comparing its expression in PCa with the adjacent normal tissue from the same patients. Another plausible reason for the discrepancy in gene expression in the paired tumor and normal samples is that the RNA abundancy for these genes does not reflect protein levels, since our IHC results show that PAGE4, NUF2 and SPAG4 are up-regulated in tumor although the mRNAs are down-regulated when compared to the normal prostate. This would also explain why for the other CTAs no differences in mRNA levels in normal and tumor contrast with significant differences at the protein level. Also, translational machinery activity and temporal mRNA and protein degradation are additional variables that can cause in discrepancies between RNA abundance and protein expression [49,50].

CTAs, especially the ones located on the X chromosome (the CT-X-Antigens), constitute a family of genes with great potential as biomarkers in different types of tumors, since they are cancer-specific and rarely expressed by normal tissues. Many of these genes when aberrantly expressed in cancer cells are immunogenic and can induce antibody- and cell-mediated responses that make them good targets for the development of cancer vaccines [24,51,52]. Unfortunately, their role as cancer biomarkers and therapeutic targets has not been appreciated in many tumor types, including PCa. The current study demonstrates that CTA expression profiles might be an important tool to predict, at the time of diagnosis, patients with higher risk to develop metastasis and that would benefit from aggressive treatments from those men with indolent disease who may have a better quality of life receiving adequate active surveillance. Here, we used small cohorts to determine CTA expression profiles; the next step is to evaluate the expression of *CEP55, NUF2, PAGE4, PBK, RQCD1, SPAG4, SSX2* and *TTK* in larger cohorts with follow-up data right from screening to the development of MPCa. Also, the development of less invasive approaches (liquid biopsies) to measure CTAs expression in circulating tumor cells, and even the presence of antibodies against these immunogenic biomarkers would be beneficial for early detection of primary tumors as well as metastasis prediction.

## 5. Conclusions

To summarize, we have demonstrated that eight CTAs are differentially expressed in PCa. The same CTAs can also be useful to discriminate locally confined tumors from metastatic tumors. These observations were detected at the gene expression and protein levels and in different patients cohorts, which provides validation of our findings across different samples and, groups of patients. CTAs are a group of genes aberrantly expressed in cancer and some present immunogenicity. Cancer specificity and immunogenicity make this class of genes unique potential biomarkers and immunotherapy targets. Here, we demonstrate that a panel of CTAs are aberrantly expressed in PCa and associated with metastatic disease suggesting their potential as biomarkers for screening and patients’ follow-up. Further studies involving broader prospective cohorts are needed to prove their usefulness as biomarkers and also the investigation of their immunogenicity is valuable and would result in new immunotherapy strategies for men with PCa.

## Acknowledgements

The authors thank the Prostate Cancer Biorepository Network for their help in obtaining samples and clinicopathologic data; the Johns Hopkins Medical Institution (JHMI) Deep Sequencing Core for performing the Nanostring approach; the Department of Pathology in JHMI for the TMAs construction and immunohistochemistry image acquisition. We further acknowledge members of the RWV.

## Author Contributions

Conceptualization, L.T.K., R.G., P.K. and R.W.V.; Development of methodology, L.T.K., S.T., T.S., P.K., R.W.; Data acquisition, L.T.K., S.T., T.S., S.M.M. R.L.V.; Analysis, L.T.K., N.M.C., S.T., T.S., S.M.M., P.K., R.W.V.; Writing, review, and/or revision of the manuscript, L.T.K., N.M.C., S.T., T.S., S.M.M., R.L.V., R.G., P.K., R.W.V.; Study supervision, L.T.K., P.K., R.W.V.

