## Supplemental files for "Cancer/Testis Antigens Differentially Expressed in Prostate Cancer: Potential New Biomarkers and Targets for Immunotherapies"

**Supplementary Table 1** – Demographic and clinicopathological characteristics of the localized and metastatic prostate cancer cohort.

| **Sample ID** | **Metastasis** | **Tissue Type** | **Age** | **PSA*** | **Stage**** | **Gleason Score**** |
| --- | --- | --- | --- | --- | --- | --- |
| L1 | No | primary prostate cancer | 64 | 19.2 | T3aN2 | 4+3 |
| L2 | No | primary prostate cancer | 63 | 8 | T2aN0 | 3+3 |
| L3 | No | primary prostate cancer | 55 | 7.5 | T3bN0 | 5+4 |
| L4 | No | primary prostate cancer | 51 | 11.7 | T2cN0 | 3+3 |
| L5 | No | primary prostate cancer | 70 | 10.8 | T3aN0 | 3+4 |
| L6 | No | primary prostate cancer | 57 | 15.3 | T3bN0 | 4+4 |
| L7 | No | primary prostate cancer | 48 | 8.2 | T3bN0 | 3+4 |
| L8 | No | primary prostate cancer | 55 | 4.4 | T2cN0 | 3+3 |
| L9 | No | primary prostate cancer | 69 | 7.58 | T2cN0 | 3+4 |
| L10 | No | primary prostate cancer | 56 | 4 | T2aN0 | 3+4 |
| L11 | No | primary prostate cancer | 75 | 2.4 | T3bN0 | 3+4 |
| L12 | No | primary prostate cancer | 60 | 7.3 | T2cN0 | 3+4 |
| L13 | No | primary prostate cancer | 65 | 4.5 | T3bN1 | 3+4 |
| L14 | No | primary prostate cancer | 59 | 14.19 | T2cN0 | 3+4 |
| L15 | No | primary prostate cancer | 52 | 4.41 | T2cNx | 3+4 |
| L16 | No | primary prostate cancer | 49 | 8 | T2cN0 | 3+4 |
| L17 | No | primary prostate cancer | 57 | 11 | T2cN0 | 3+4 |
| L18 | No | primary prostate cancer | 64 | 6.01 | T2cN0 | 3+4 |
| L19 | No | primary prostate cancer | 58 | 4.3 | T3aN0 | 4+3+5 |
| L20 | No | primary prostate cancer | 57 | 6.1 | T2cNx | 3+3 |
| M1 | Yes | lymph node met | 86 | 1508.4 | - | - |
| M2 | Yes | liver met | 64 | 182.7 | - | - |
| M3 | Yes | bladder met | 67 | 0.72 | - | - |
| M4 | Yes | lymph node met | 90 | 1455.9 | - | - |
| M5 | Yes | liver met | 83 | 114.83 | - | - |
| M6 | Yes | lymph node met | 81 | 1276 | - | - |
| M7 | Yes | liver met | 81 | 122.76 | - | - |
| M8 | Yes | liver met | 69 | 96.21 | - | - |
| M9 | Yes | liver met | 60 | 620.5 | - | - |
| M10 | Yes | lymph node met | 70 | 2017.6 | - | - |
| M11 | Yes | lymph node met | 72 | 298.7 | - | - |
| M12 | Yes | lymph node met | 70 | 1493.2 | - | - |
| M13 | Yes | lymph node met | 84 | 240.65 | - | - |
| M14 | Yes | lymph node met | 73 | 55.58 | - | - |
| M15 | Yes | adrenal met | 71 | 735.97 | - | - |
| M16 | Yes | lung met | 82 | 2295 | - | - |
| M17 | Yes | lymph node met | 64 | 3869 | - | - |
| M18 | Yes | pancreas met | 76 | 5690.8 | - | - |
| M19 | Yes | lymph node met | 77 | 3700 | - | - |
| M20 | Yes | kidney met | 62 | 967.48 | - | - |

* PSA levels correspond to the last test result available before tissue samples were collected.

** Information regarding tumor stage and Gleason score were not available for the metastatic prostate cancer (MPCa) cases. Samples were collected during autopsy from patients diagnosed with MPCa with previous radical prostatectomy.

**Supplementary Table 2 –** Clinicopathological information available for the paired prostate cancer and benign adjacent cohort.

| **Sample ID** | **Gleason Score** | **P Stage** |
| --- | --- | --- |
| PT1 and PN1 | 3+4 | T2N0Mx |
| PT2 and PN2 | 3+3 | T2N0Mx |
| PT3 and PN3 | 3+3 | T2N0Mx |
| PT4 and PN4 | 3+4 | T2N0Mx |
| PT5 and PN5 | 3+4 | T2N0Mx |
| PT6 and PN6 | 3+4 | T2N0Mx |
| PT7 and PN7 | 3+4 | T2N0Mx |
| PT8 and PN8 | 4+4 | T3BN1Mx |
| PT9 and PN9 | 4+5 | T3AN1Mx |
| PT10 and PN10 | 4+4 | T3AN0Mx |
| PT11 and PN11 | 4+5 | T3AN0Mx |
| PT12 and PN12 | 3+3 | T2XN0Mx |
| PT13 and PN13 | 4+4 | T3AN0Mx |
| PT14 and PN14 | 3+3 | T2N0Mx |
| PT15 and PN15 | 3+3 | T2N0Mx |
| PT16 and PN16 | 4+3 | T3AN0Mx |
| PT17 and PN17 | 4+3 | T3AN0Mx |
| PT18 and PN18 | 4+3 | T3AN0Mx |
| PT19 and PN19 | 4+3+5 | T3AN0Mx |
| PT20 and PN20 | 4+4 | T3BN1Mx |
| PT21 and PN21 | 4+5 | T3AN0Mx |
| PT22 and PN22 | 4+5 | T3AN1Mx |
| PT23 and PN23 | 4+3+5 | T3AN0Mx |
| PT24 and PN24 | 4+5 | T3BN1Mx |
| PT25 and PN25 | 4+4+3 | T3AN0Mx |

**Supplementary Table 3 –** Prostate cancer patients’ demographics in TMA 681 and 682 (N=80).

| **Variable** | **Recurrence** | | **Total** |
| --- | --- | --- | --- |
|  | **No** | **Yes** |  |
| Age | 59.04±6.57 | 58.82±5.80 | - |
| Gleason score |  |  |  |
| 6 | 7 | 1 | 8 |
| 7 (3+4) | 14 | 3 | 17 |
| 7 (4+3) | 15 | 3 | 18 |
| ≥8 | 11 | 15 | 26 |
| TNM stage |  |  |  |
| T2 | 26 | 2 | 28 |
| T3A | 16 | 15 | 31 |
| T3B | 3 | 5 | 8 |
| Gland weight (g) | 51.94±14.90 | 56.19±21.38 | - |
| PSA (ng/mL) | 8.29±5.29 | 13.15±10.15 | - |
| PSAD (ng/mL/g) | 0.17±0.11 | 0.25±0.20 | - |
| Race |  |  |  |
| African American | 6 | 3 | 9 |
| Hispanic | 1 | 0 | 1 |
| Caucasian | 37 | 18 | 55 |
| Others | 3 | 1 | 4 |
| Surgical Margin |  |  |  |
| Negative | 41 | 16 | 57 |
| Positive | 6 | 6 | 12 |
| Seminal vesicle status |  |  |  |
| Negative | 46 | 16 | 62 |
| Positive | 1 | 6 | 7 |
| Lymph node status |  |  |  |
| Negative | 46 | 22 | 68 |
| Positive | 1 | 0 | 1 |
| Capsular penetration (fcp) |  |  |  |
| Negative | 38 | 16 | 54 |
| Positive | 8 | 6 | 14 |
| Capsular penetration (ecp) |  |  |  |
| Negative | 34 | 8 | 42 |
| Positive | 8 | 6 | 14 |
| Organ confinement status |  |  |  |
| Non-confined | 21 | 20 | 41 |
| Confined | 26 | 2 | 28 |

1. Age, gland weight, PSA and PSAD data are shown as mean ± SD.

2. PSAD (ng/mL/g) = PSA (ng/mL)/gland weight (g)

**Supplementary Table 4 –** qRT-PCR primers’ sequences used for CTA gene expression analysis.

| **Gene** | **Forward primer (5'-3')** | **Reverse primer (5'-3')** |
| --- | --- | --- |
| *ACTB* | CCTGGCACCCAGCACAAT | GCCGATCCACACGGAGTACT |
| *CEP55* | ACTGTGGCTCCAAACTGCTT | GAGCAGCTGTTTCCGTTTTC |
| *CSAG2* | AGTGGGCCAACACTATCCAG | CTGGCTGTCCGAAGAGAGAC |
| *CTAG1B* | GACACCCATGGAAGCAGAG | GACAGGAGCTGATGGAGAGC |
| *JARID1B* | CTTCTTGTTTGCCTGCATCA | ATTTTGGGATTTCCCTCCAC |
| *MAGEA1* | GCCTTTCCCACTACCATCAA | GGAGCAGAAAACCAACCAAA |
| *MAGEA2* | CAGCAACCAAGAAGAGGAGG | TGCAAGTACTCGGAGGCTTT |
| *MAGEA6* | AGTAGGAAGGTGGCCGAGTT | CAAGGAACTGGAAGCTTTGC |
| *MAGEA12* | CTGGAGTCAATCCGATGAGG | TCTCCAGGTCAGGAAAGGTG |
| *NOL4* | CAGCCACTGAACCTGAGTGA | GCCTCTCGTTCCATCTTGAG |
| *NUF2* | TGCCGTGAAACGTATATGGA | ATTAATGCCTCCTGGTGTGC |
| *PAGE4* | CGTAAAGTAGAAGGTGATTG | ATGCTTAGGATTAGGTGGAG |
| *PBK* | TCTCATTCTCCTTGGGCTGT | TGCCATCATTGGCTTCAGTA |
| *PLAC1* | TGCTGTGCTCCATAGACTGG | GTGTGGCTGAACATGGTTTG |
| *RQCD1* | GCGAGAATCTGTTCCTGACC | GGTGGGTGGGTTGATAGATG |
| *SEMG1* | AAACCTCACTCTGTCCTGCG | TCGGCCATGCTGTTGATCTT |
| *SPAG4* | ATGAGGATTTTGTGCGGAAG | GCGTAATCGTGGGATGTCTT |
| *SSX2* | GGTGGAGCAGTCAGAACACA | TGGGTCCCTGTTGTGTGTAA |
| *SSX4* | AGGAGACCCAGGGATGATG | GTCATGACCTCATAGTTTAGCTTCA |
| *TMEFF2* | TGCATGGGAAGTGTGAGCAT | CAGGACCGGGAACAACGTAT |
| *TMEM108* | AGACACTGGGATGGTCCTTG | GCAAAGTGATGCCTGCTGTA |
| *TPTE* | TTGGAGAAAGGCGAACAGAT | TTGGAGGGAGATTCCAGTTG |
| *TTK* | CAGCAGCAACAGCATCAAAT | TGCTTGAACCTCCACTTCCT |

**Supplementary Table 5 –** Immunohistochemistry primary antibodies information.

| **Protein** | **Dilution** | **Manufacturer** | **Catalog #** |
| --- | --- | --- | --- |
| CEP55 | 1:2000 | Abcam | ab170414 |
| NUF2 | 1:75 | Sigma-Aldrich | SAB2700437 |
| PAGE4 | 1:1000 | Sigma-Aldrich | HPA023880 |
| PBK | 1:200 | Sigma-Aldrich | HPA005753 |
| RQCD1 | 1:10 | Sigma-Aldrich | HPA046622 |
| SPAG4 | 1:300 | Abcam | ab183106 |
| SSX2 | 1:50 | Abcam | ab117972 |
| TTK | 1:250 | Sigma-Aldrich | HPA016834 |

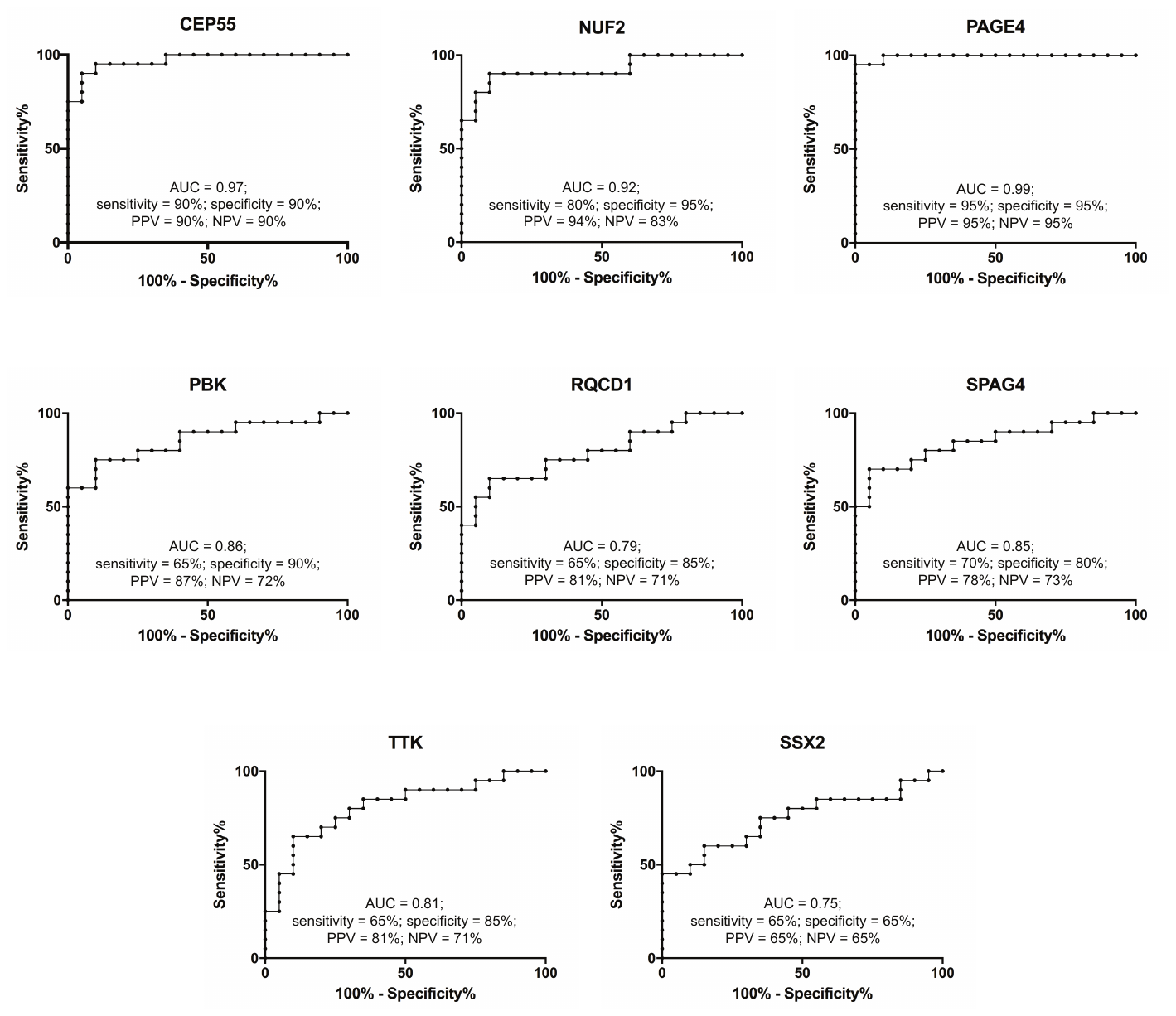

**Supplementary Figure 1 –** Receiver operating characteristic (ROC) curve analysis for the Nanostring gene expression analysis. In the figure are represented the ROC curves for the CTAs with area under curve (AUC) above the cutoff (0.7). For each CTA sensitivity, specificity, positive (PPV) and negative (NPV) predictive values are also shown.

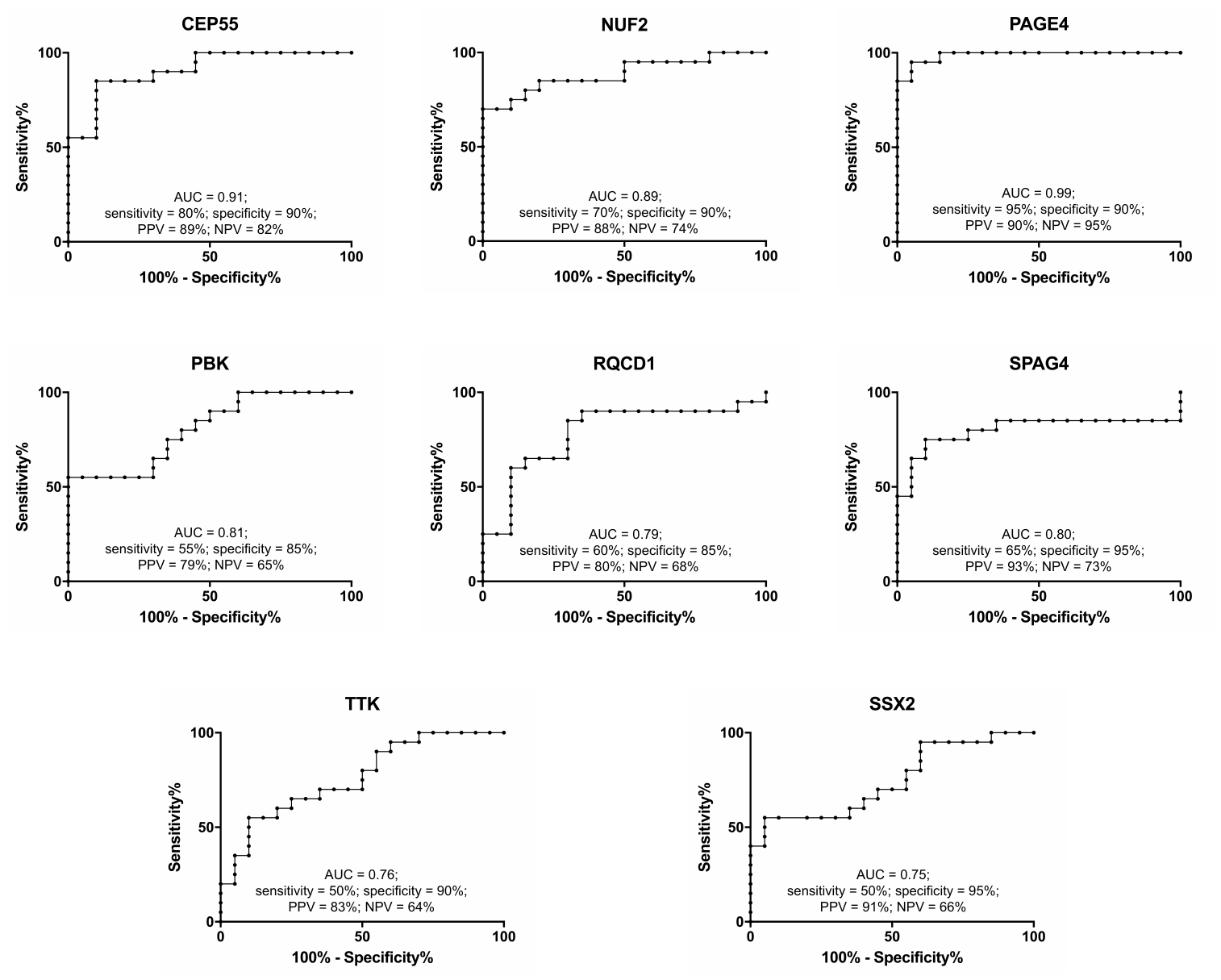

**Supplementary Figure 2 –** Receiver operating characteristic (ROC) curve analysis for the qRT-PCR gene expression analysis (Nanostring technical validation). In the figure are represented the ROC curves for the CTAs with area under curve (AUC) above the cutoff (0.7). For each CTA sensitivity, specificity, positive (PPV) and negative (NPV) predictive values are also shown.
